# Juicebox.js provides a cloud-based visualization system for Hi-C data

**DOI:** 10.1101/205740

**Authors:** James Robinson, Douglass Turner, Neva C. Durand, Helga Thorvaldsdottir, Jill P. Mesirov, Erez Lieberman Aiden

## Abstract

*Contact mapping experiments such as Hi-C explore how genomes fold in 3D. Here, we introduce Juicebox.js, a cloud-based web application for exploring the resulting datasets. Like the original Juicebox application, Juicebox.js allows users to zoom in and out of such datasets using an interface similar to Google Earth. Furthermore, Juicebox.js encodes the exact state of the browser in a shareable URL. Creating a public browser for a new Hi-C dataset does not require coding and can be accomplished in under a minute*.

## MAIN TEXT

Hi-C and other contact mapping experiments measure the frequency of physical contact between loci in the genome. The resulting dataset, called a “contact map”, is often represented using a two-dimensional heatmap where the intensity of each pixel indicates the frequency of contact between a pair of loci. The highest resolution Hi-C heatmaps presently available contain trillions of pixels and exhibit structures across a wide range of size scales. To explore such data, we recently developed Juicebox (Durand, Robinson, et al., 2016), a desktop application inspired by Google Earth. Juicebox enables users to interactively zoom in and out of Hi-C datasets and to perform many other functions.

One important use case for any scientific data visualization system is to enable multiple research groups to easily recapitulate a finding, thereby ensuring greater reproducibility of the scientific record. A web-based application enabling the visualization of Hi-C contact maps would make it easier to communicate findings as compared to a desktop application. Although several web-based Hi-C contact map browsers exist (Lieberman-Aiden, van Berkum, et al., 2009, Wang, et al., 2017, Yardimici, et al., 2017) none of the tools that have been published to date support interactive zooming in real-time, a critical feature that is essential to seamless exploration of Hi-C data (though see (Kerpedjiev, et al., 2017)).

Here we present Juicebox.js, a web application that implements many core features of the Juicebox desktop application (Figure 1). Juicebox.js enables users to load one or more contact maps, to zoom in and out, and to compare the contact maps to genomic tracks and 2D annotations. Juicebox.js supports both desktop and mobile devices; users can explore the maps using a keyboard and mouse, or by means of touch-screen gestures (such as pinch-zoom).

**Figure 1.**
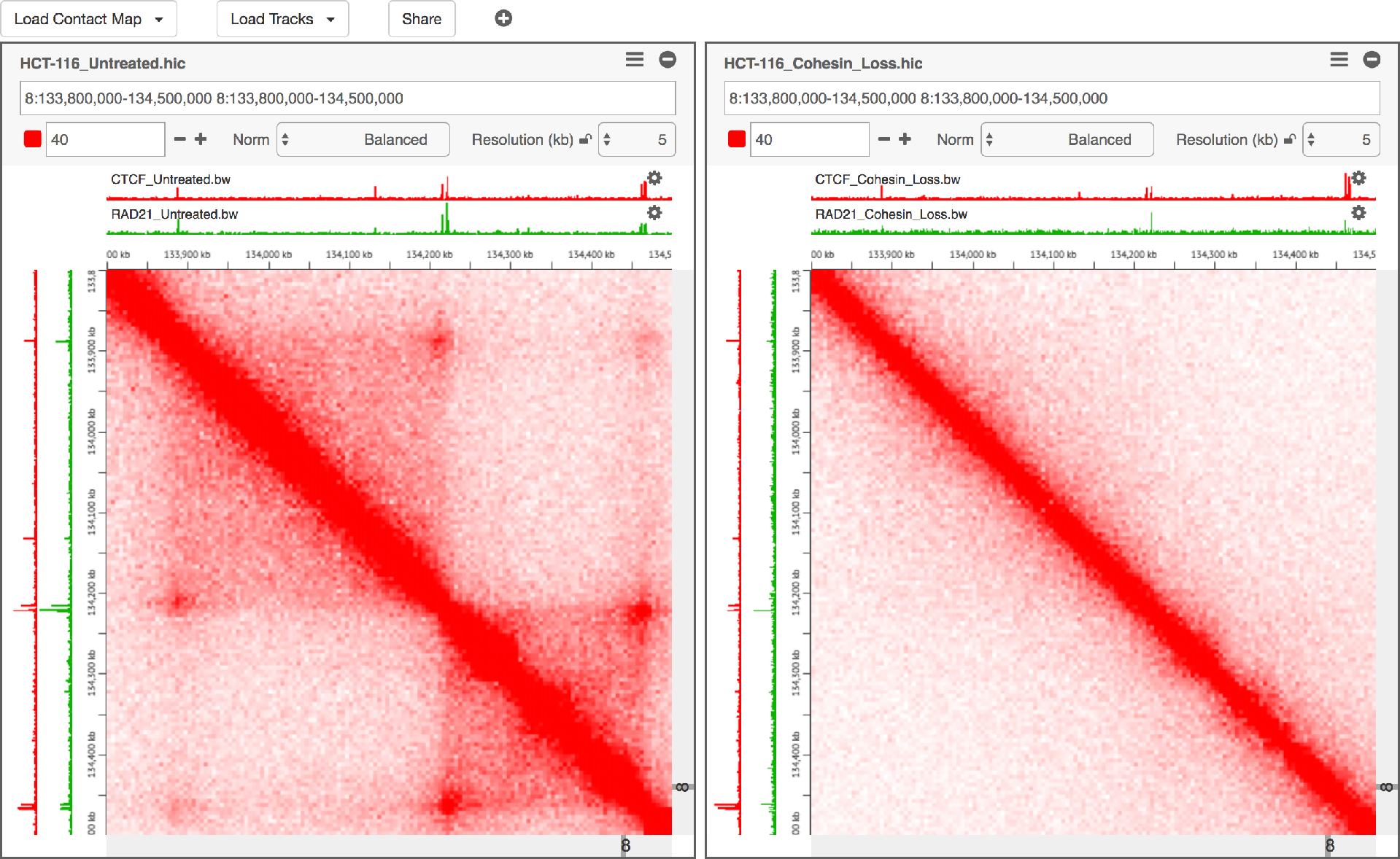
Juicebox.js makes it easy to share interactive visualizations of contact mapping data derived from Hi-C and other experiments. Hi-C maps from new experiments can be easily added and juxtaposed with tracks from ENCODE and other sources. It is possible to zoom in and out in real-time using either a mouse or touch-screen gestures. Display parameters like the color scale and the normalization can be adjusted interactively. The complete state of the browser can always be encoded as a sharable URL. No programming is necessary to share and explore new datasets. *Left:* A loop resolution Hi-C map showing all contacts within an 700kb genomic interval, generated using HCT-116 human colorectal carcinoma cells. Loops, which form here due to physical tethering between two CTCF- and cohesin-bound loci, manifest as bright peaks away from the diagonal. Contact domains, genomic intervals that exhibit enhanced contact frequency within themselves, manifest as bright squares along the diagonal. When the two anchors of a loop demarcate a contact domain, the resulting feature is called a "loop domain". Right: The same region in HCT-116 cells after the RAD21 subunit of the cohesin complex has been degraded using an auxin-inducible degron system. The loop domains all disappear completely, demonstrating that they are dependent on cohesin. An interactive version of this figure is available at http://www.cell.com/cell/9802-figure-2.

Juicebox.js is designed to work with data in the *hic* format (Durand, Robinson, et al., 2016), a compressed indexed format that enables fast query and can be used with a wide variety of contact mapping experiments, such as dilution Hi-C (Lieberman-Aiden, van Berkum, et al., 2009), *in situ* Hi-C (Rao, Huntley, et al., 2014), single-cell Hi-C (Nagano, et al., 2013), Hi-C^2^ (Sanborn, Rao, et al., 2015), ChIA-PET (Fullwood, et al., 2009), and HiChIP (Mumbach, et al., 2016). This format has been adopted by many groups and several large consortia, including the Encyclopedia of DNA Elements (ENCODE) and the 4D Nucleome Consortium.

Users of juicebox.js can load a *hic* file from their local hard-disk or file system. However, because *hic* files encode billions of contacts, they can be very large (hundreds of gigabytes) and keeping local copies is often unwieldy. To address this need, juicebox.js can access *hic* files located at an arbitrary URL, downloading only the small portion of the file that is required to fulfill each user request. Users can also remotely access genomic tracks (such as ChIP-Seq data) and 2D annotations (such as Hi-C loop calls) in all standard formats (*BigWig, bed, bedpe*, etc.) via URL. Loading via URL works with any standard file server, and with a range of cloud storage providers such as Amazon S3, Dropbox, and Google Drive. As an example, Juicebox.js is designed to automatically connect to ENCODE servers and remotely load any dataset generated by the consortium.

Juicebox.js has many features to facilitate reproducibility and sharing.

First, the complete state of any Juicebox.js instance can be encoded in a sharable URL link. The exact same fully-interactive state can be recapitulated merely by opening this URL in another browser (including mobile browsers). The URL can also be shared with the scientific public in a journal article, using social media such as Twitter, or even by means of a QR code. Sharable URLs can be created directly from Juicebox or programmatically by creating a URL text string with the necessary parameters included (such as the files to be accessed, genomic location, and color scale). For instance, a user could write a script to create a large number of URLs corresponding to a large number of features that have been identified in a given Hi-C dataset. Any of these features could then be explored further simply by clicking the corresponding link.

Second, Juicebox.js instances, in any desired state, can be embedded in a webpage using a few lines of code. This makes it simple to include Juicebox.js on a blog, in a news article, or in an online journal article. By embedding one or more Juicebox.js instances on a single webpage, it is possible to take a static publication figure and create an online version in which all contact maps are fully interactive. This enables other researchers to explore the results by changing the color scale and other display properties, and to ensure that an image is representative of the dataset as a whole, rather than a cherry-picked example. For a recent publication, we used Juicebox.js to create 5 interactive figures, which the journal was able to host alongside the online version of the paper (Rao, et al., 2017).

Finally, Juicebox.js is a purely client-side application: it reads data directly from files hosted on standard webservers, but no application code is required on the server. Consequently, Juicebox.js can easily support an almost unlimited number of users. This also makes it possible to create interactive browsers with Juicebox.js for one or more *hic* files without writing a single line of code. For instance, a user can simply upload a *hic* file to Dropbox; use Dropbox to create a link to the file; and then load that link into Juicebox.js. The resulting Juicebox.js URL can immediately be shared with users all over the world, enabling them to interactively explore the data located on the Dropbox folder.

An instance of Juicebox.js is available at aidenlab.org/juicebox. The code, which is available at github.com/igvteam/juicebox.js, is open source, and is licensed under the MIT license. The test procedure and datasets associated with this publication are available at: https://data.mendeley.com/archiver/fbgc85km6j. We hope that Juicebox.js will greatly facilitate the sharing of Hi-C and other contact mapping datasets.

## ACKNOWLEDGMENTS

We thank Suhas Rao and Sheikh Russell for assistance creating the figure. This work was supported by an NIH New Innovator Award (1DP2OD008540-01), an NSF Physics Frontiers Center Award (PHY-1427654, Center for Theoretical Biological Physics), the Welch Foundation (Q-1866), an NVIDIA Research Center Award, an IBM University Challenge Award, a Google Research Award, a Cancer Prevention Research Institute of Texas Scholar Award (R1304), a McNair Medical Institute Scholar Award, an NIH 4D Nucleome Grant U01HL130010, an NIH Encyclopedia of DNA Elements (ENCODE) Mapping Center Award UM1HG009375, the President’s Early Career Award in Science and Engineering to E.L. A., an NIH National Cancer Institute (NCI) Award (R01CA15730), and an NIH NCI Informatics Technology for Cancer Research (ITCR) Award (U24CA210004)

## AUTHOR CONTRIBUTIONS

E.L.A. and J.T.R. conceived of this project. J.T.R., D.T., N.C.D., and H.T. created the tool. J.P.M. contributed to tool development. J.T.R. and E.L.A. wrote the manuscript.

